# Terrestrial Support of Aquatic Food Webs via an Overlooked Pathway—Inorganic Carbon and its Significance to Global Carbon Cycle

**DOI:** 10.1101/2023.03.27.534453

**Authors:** Junna Wang, John R. Durand, Sharon P. Lawler, Pinyuan Chen, Xiaoli Dong

## Abstract

Freshwater ecosystems receive substantial terrestrial organic matter (t-OM) from surrounding landscapes. How the t-OM is transferred affects aquatic food webs and global carbon budgets. Previous studies have emphasized terrestrial support of aquatic ecosystems via direct organic carbon subsidy, overlooking the dissolved inorganic carbon (DIC) pathway, that is, DIC from t-OM decomposition is used by aquatic primary producers, supporting higher trophic levels. Using 2-year ^13^C and ^15^N measurements of phytoplankton, zooplankton, terrestrial plants, sediments, dissolved and particulate organic matter from seasonal wetlands, we found that while zooplankton (mid-trophic consumers) used t-OM directly in January, in March and May zooplankton were mainly supported by phytoplankton that used DIC recycled from t-OM mineralization and methanogenesis. The dominance of this DIC pathway is tightly coupled with the characteristics of these systems. Mineralization and methanogenesis of rich fresh t-OM resulted in supersaturated CO_2_ with high CO_2_ and CH_4_ emissions. Atmospheric CO_2_ diffusion and methanogenesis significantly enriched *δ*^13^*C* of DIC, leading to wide variations in *δ*^13^*C* of DIC between -12.4 and 6.7 ‰, which provided ideal conditions to quantify carbon cycling in these widespread but understudied ecosystems. Our findings draw attention to potentially high carbon emissions from temporary freshwater ecosystems that are being increasingly common under warming climate.

## Introduction

Terrestrial organic matter (t-OM) delivered to aquatic ecosystems (allochthonous resources) and organic matter produced within aquatic ecosystems (autochthonous resources) are two distinct resources supporting aquatic food webs (Pace et al. 2004) (Fig. 1A). Compared to t-OM, autochthonous OM is often higher in nutrient content and more easily assimilated, and has been thought to be the primary resource supporting aquatic ecosystems (Brett et al. 2009). Over the last two decades, evidence from ^13^C addition experiments, stable isotope (*δ*^13^*C, δ*^15^*N, δ*^2^*H*) analysis, and feeding experiments suggests substantial t-OM subsidies to aquatic food webs (Carpenter et al. 2005; Taipale et al. 2008; Cole et al. 2011; Harfmann et al. 2019). While efforts have been made to quantify and explain the varying contributions of these two resources (Tanentzap et al. 2017), the two resources are not independent. Dissolved inorganic carbon (DIC) used for autochthonous primary production can be from t-OM decomposition—an inorganic carbon pathway of terrestrial subsidy that has been largely overlooked (Lennon et al. 2006; Demars et al. 2020). Such a neglect may be partly due to the assumption that primary producers use DIC mainly from atmospheric carbon dioxide (CO_2_), but accumulating studies show that most freshwater ecosystems are net carbon sources (Santos et al. 2022).

**Figure 1.**
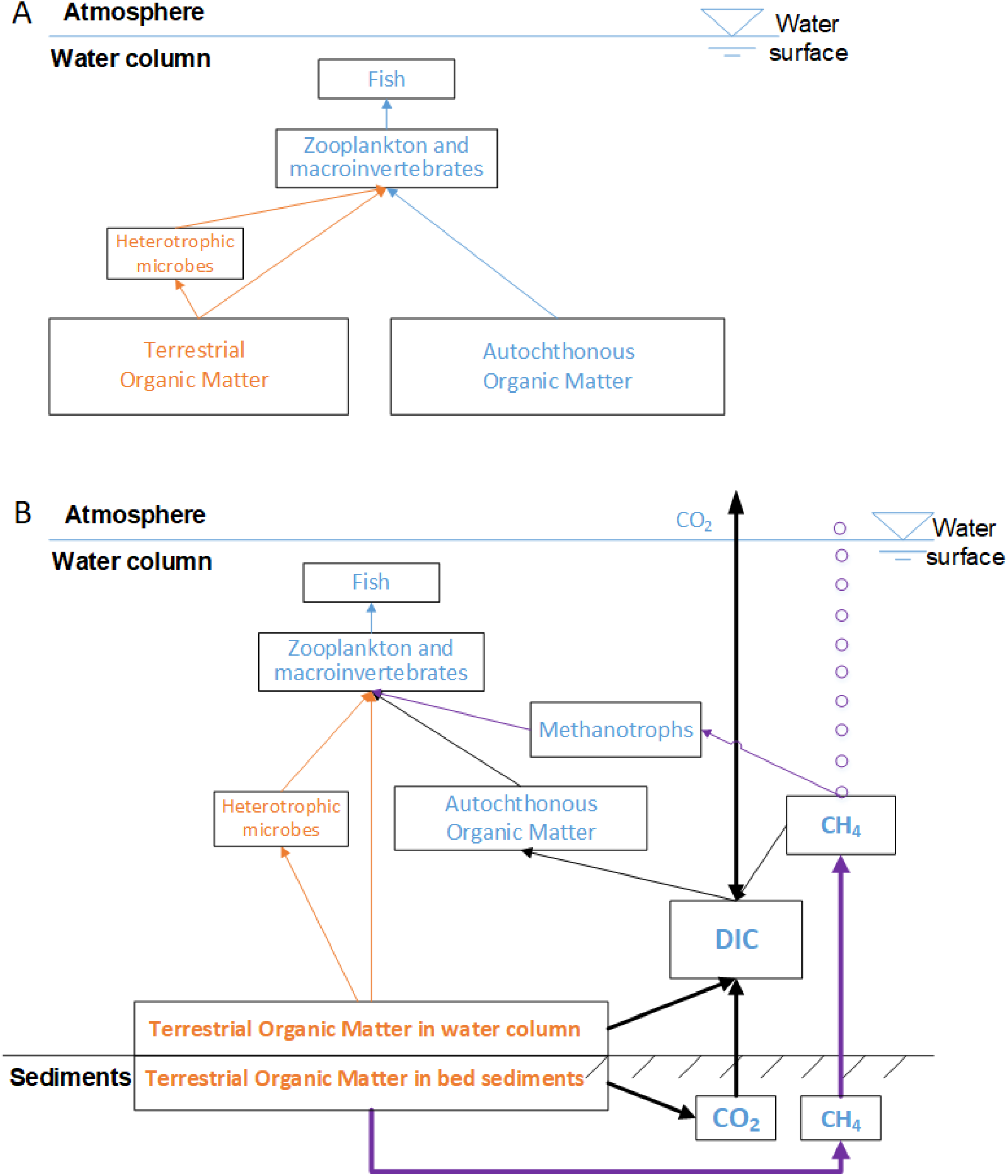
Conventional perspective on terrestrial support of aquatic ecosystems (A) and proposed (B) perspective on terrestrial support of small, shallow, lentic freshwater ecosystems with rich terrestrial detritus inputs. DOC: dissolved organic carbon; POM: suspended particulate organic matter; SOM: sediment organic matter; and DIC: dissolved inorganic carbon. In (B), orange arrows represent the organic carbon pathways, black arrows represent inorganic carbon pathways, and purple arrows represent the methane pathway. Arrow thickness qualitatively represents the relative size of each flux.

We propose a new perspective on terrestrial subsidy of aquatic ecosystems as occurring via three pathways (Fig. 1B): organic carbon (OC, orange arrows), DIC (black arrows), and methane (CH_4_, purple arrows). This perspective emphasizes the importance of the DIC pathway for the global carbon cycle and aquatic food webs. In view of resolving global carbon budgets, if autochthonous primary producers mainly use DIC recycled from decomposing t-OM instead of atmospheric CO_2_, carbon sequestration of aquatic ecosystems may be lower than expected (Webb et al. 2019), and CO_2_ emission from aquatic ecosystems may be enhanced with increasing t-OM inputs (Lapierre et al. 2013). If DIC used by primary producers is mainly produced by anaerobic methanogenesis, the co-product, methane (CH_4_), likely makes certain aquatic ecosystems CH_4_ emission hotspots (Bastviken et al. 2011). Furthermore, CH_4_ can be incorporated into aquatic food webs via consumption of methane-oxidizing bacteria—methanotrophs by consumers (Stanley et al. 2016), forming the CH_4_ pathway (Fig. 1B). In contrast to the conventional framework describing the two resources for aquatic food webs, t-OM and autochthonous OM separately (Fig. 1A), we emphasize the linkage between the two and highlight the multiple pathways through which t-OM supports aquatic food webs, and its role in maintaining stability and productivity of aquatic ecosystems (Moore et al. 2004).

Here, we used aquatic ecosystems in seasonal wetlands to investigate terrestrial support of freshwater ecosystems via multiple pathways, and its implication for the global carbon cycle. Our system represents widespread but under-studied small, shallow, temporary, aquatic ecosystems that feature large fresh t-OM inputs and long residence time. We first characterized *δ*^13^*C* signatures in DIC, dissolved organic carbon (DOC), suspended particulate organic matter (POM), phytoplankton, terrestrial plants, and zooplankton. We then quantified the contribution of t-OM to DIC and emission rates of CO_2_ and CH_4_ using mass balance models of DIC and DI^13^C. Lastly, we used Bayesian mixing models to estimate relative contributions of different pathways in support of aquatic food webs.

## Methods

### Study sites

Our sites include five seasonal wetlands and one rice field located on a floodplain adjacent to the Sacramento River (Fig. S1A). The region features a Mediterranean climate with wet winters and dry summers. Around 84% of annual precipitation (∼498 mm) occurs in November-March. These sites are filled with rainfall or pumped water in November-December to a water depth < 0.5 m to mimic historical flooding. Water levels do not vary much before March, but decline rapidly afterwards due to little precipitation, rising temperature and evaporation. The rice field is drained in March, and wetlands dry naturally in April-June. During the aquatic phase, wetland plants are restricted to edges and scattered islands (Fig. S1B). In the dry phase, they occupy the entire wetlands.

### Sample collection and stable isotope analysis

We conducted six sampling events in January, March, and May of 2021 and 2022 (twice in May, 2021, none in January, 2022; Table S1). In each sampling event, we collected water samples and solid samples including surface sediments, phytoplankton, zooplankton, and terrestrial plants that represent t-OM at 2∼3 replicated locations with water depth <0.3 m. We measured water depth and in-situ water quality including temperature, pH, Chlorophyll-a (Chla), and dissolved oxygen. Water samples were analyzed for DIC, DOC, and total organic carbon (TOC), *δ*^13^*C* of DOC, *δ*^13^*C* of DIC, and *δ*^13^*C* of POM. We washed all solid samples except sediments, dried and ground them to measure their organic *δ*^13^*C* and *δ*^15^*N*. Analytical details are provided in SI text S1.

### Mass balance models for DIC and DI^13^C

To quantify the contribution of t-OM to DIC and emission rates of CO_2_ and CH_4_, we constructed mass balance models for DIC and DI^13^C. DIC in our systems is consumed by aquatic photosynthesis, and produced by mineralization of DOC and POC, anaerobic methanogenesis of sediment organic carbon (SOC), and oxidation of CH_4_ (Fig. S2) (Bade et al. 2004). It is also affected by air-water exchange between atmospheric and aqueous CO_2_ (CO_2_(aq)) (Himmelblau 1964). The effect of carbonate weathering in groundwater inputs is negligible because groundwater levels are > 20 m below land surface (USGS: 382515121214401). The DIC-related processes (Fig. S2) carry distinct *δ*^13^*C* signatures. DIC produced by mineralization has a *δ*^13^*C* signature close to source carbon. Due to similarity of *δ*^13^*C* in DOC and POC (see Results), we lumped DOC and POC, and used average *δ*^13^*C* as *δ*^13^*C* of TOC. SOC methanogenesis produced equal moles of CH_4_ and CO_2_, but CH_4_ carries very depleted *δ*^13^*C* (−80∼-42 *‰*), while CO_2_ has extremely enriched *δ*^13^*C* (0∼20 *‰*) (Gu et al. 2004). Of CH_4_ produced by methanogenesis, more than half is emitted to the atmosphere via ebullition and diffusion (Baulch et al. 2011). The remainder is oxidized to CO_2_ in water column with a *δ*^13^*C* signature ∼10 *‰* depleted relative to source CH_4_ (Shoemaker and Schrag 2010). Therefore, methanogenesis can significantly enrich *δ*^13^*C* of DIC if most CH_4_ is emitted (Gu et al. 2004). Aquatic photosynthesis can also enrich *δ*^13^*C* of DIC because algae selectively take up CO_2_(aq) with depleted *δ*^13^*C*, when CO_2_ is not limiting (Hollander and McKenzie 1991). Air-water CO_2_ exchange can enrich or deplete *δ*^13^*C* of DIC depending on *δ*^13^*C* of DIC relative to atmospheric equilibrium *δ*^13^*C* (δ^13^C_DIC_(eq)). Mass balance models were constructed for each sampling event at each site and are expressed as:

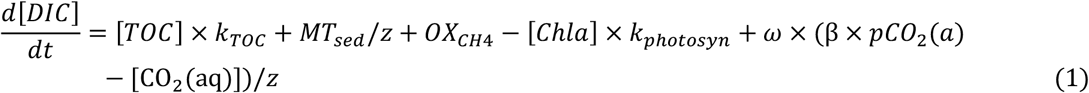

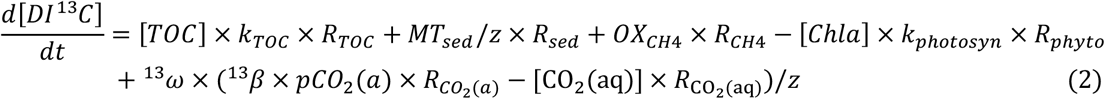

where [DIC], [TOC], [CO_2_(aq)], and [DI^13^C] denote concentrations of DIC, TOC, CO_2_(aq), and DI^13^C (mol m^-3^); [Chla] is concentration of Chla (mg m^-3^); *t* is time (d); *k*_*TOC*_ is TOC mineralization rate (1/d); *MT*_*sed*_ is CO_2_ production rate by methanogenesis (mol m^-2^ d^-1^); *z* is water depth (m); *OX*_*CH*4_ is methane oxidation rate (mol m^-3^ d^-1^); *k*_*photosyn*_ is DIC uptake rate per unit Chla (mol mg^-1^ d^-1^); *ω* and ^13^*ω* are gas exchange coefficients of CO_2_ and ^13^CO_2_ (m d^-1^); *β* and ^13^*β* are solubility of CO_2_ and ^13^CO_2_ (mol m^-3^ atm^-1^) (Wanninkhof 1985); *pCO*_2_(*a*) is partial pressure of CO_2_ in the air (atm); 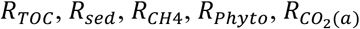, and 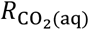 represent ^13^*C*/^12^*C* ratios (unitless) of TOC, DIC released by methanogenesis, DIC produced by methane oxidation, DIC used by phytoplankton, atmospheric CO_2_, and CO_2_(aq). Details about calculating these ^13^*C*/^12^*C* ratios, *ω*, ^13^*ω, β*, ^13^*β*, and [CO_2_(aq)] are provided in SI text S2. We assumed that 30% of CH_4_ produced by methanogenesis was oxidized in water column (i.e., *OX*_*CH*4_ = 0.3 × *MT*_*sed*_ /*z*), and the remaining was emitted (McNicol et al. 2020). We estimated the unknown parameters: *k*_*TOC*_, *MT*_*sed*_, and *k*_*photosyn*_ by Bayesian statistical inference.

### Bayesian mixing models

To estimate relative strengths of the three pathways (Fig. 1B), we constructed Bayesian mixing models of *δ*^13^*C* and *δ*^15^*N* for Daphnia and Copepoda, two dominant zooplankton taxa, for each sampling event at each site (Fig. S3). Since heterotrophic microbes decomposing t-OM have the same isotopic signatures as t-OM (Shoemaker and Schrag 2010), we used t-OM’s isotope signatures to represent isotope signatures of the OC pathway. At our sites, t-OM includes C3 and C4 plants with distinct *δ*^13^*C* signatures (Fig. S4). We separated them in the models:

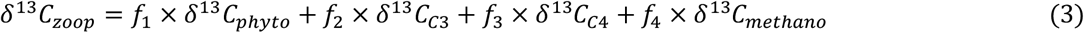

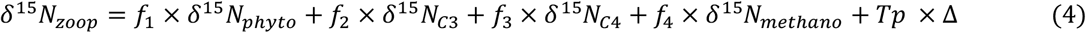

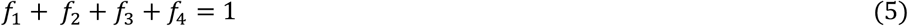

where *δ*^13^*C*_*zoop*_, *δ*^15^*N*_*zoop*_, *δ*^13^*C*_*phyto*_, *δ*^15^*N*_*phyto*_, *δ*^13^*C*_*C*3_, *δ*^15^*N*_*C*3_, *δ*^13^*C*_*C*4_, *δ*^15^*N*_*C*4_, *δ*^13^*C*_*methano*_, and *δ*^15^*N*_*methano*_ are *δ*^13^*C* and *δ*^15^*N* of zooplankton, phytoplankton, C3 plants, C4 plants, and methanotrophs (‰); *f*_1_, *f*_2_, *f*_3_, and *f*_4_ are relative contributions of the four resources to zooplankton biomass (unitless); *Tp* is trophic level of zooplankton (unitless); and Δ is per trophic level *δ*^15^*N* enrichment (‰). We measured *δ*^13^*C* and *δ*^15^*N* of zooplankton, phytoplankton, C3 and C4 plants. We set Δ = 2.52 ‰, *δ* ^13^*C*_*methano*_ = −68 ± 10 ‰, and *δ* ^15^*N*_*methano*_ = 1.8 ± 1 ‰ (Taipale et al. 2008; Cole et al. 2011; Solomon et al. 2011). We inferred values of *f*_1_, *f*_2_, *f*_3_, *f*_4_, and *Tp* using Bayesian inference with 95% credible interval.

## Results

### Aquatic ecosystems in seasonal wetlands as continuous CO_2_ sources

Aqueous CO_2_ was always supersaturated across all six sites during our sampling periods. CO_2_ supersaturation level varied between 171% and 2,153%, and CO_2_ emission rates varied between 0.059 and 1.678 gC m^-2^ d^-1^ (Fig. 2A; Table S1), suggesting that these aquatic ecosystems were likely continuous CO_2_ sources during their lifespans. Partial pressure of aqueous CO_2_ (*p*CO_2(aq)_) increased slightly from January to March, but decreased significantly from March to May (*t* = -3.44, *df* = 5, *p* = 0.02). *p*CO_2(aq)_ varied greatly among sites, with the highest recorded *p*CO_2(aq)_ and CO_2_ emissions both occurring in the rice field (Fig. 2A; Table S1).

**Figure 2.**
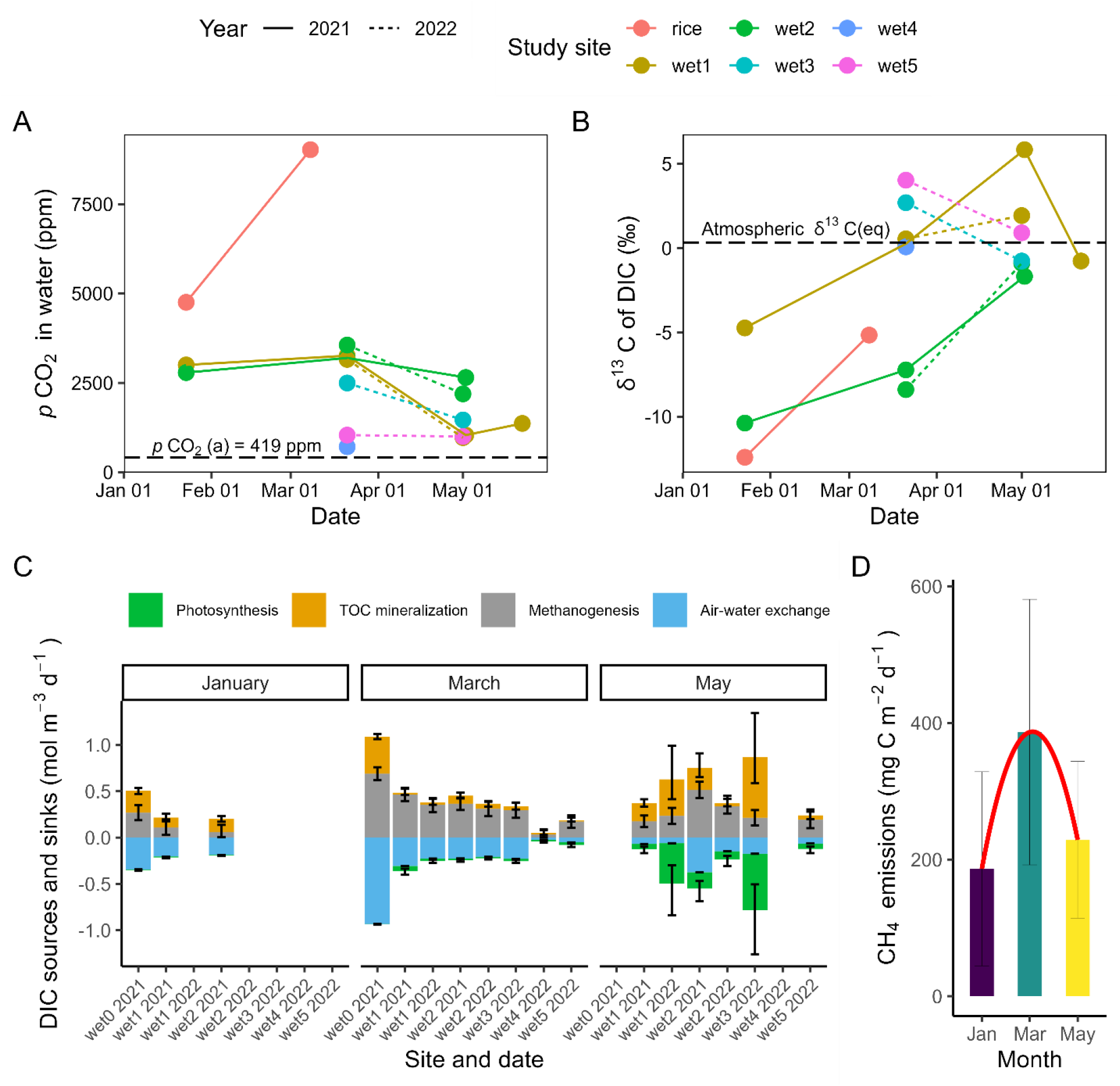
Temporal variations in *p*CO_2_ in water column (A), in *δ* ^13^*C* of DIC (B) in the rice field (“rice”) and the five wetlands (“wet1”∼”wet5”) in 2021 and 2022, monthly sources (+) and sinks (−) of DIC (C), and the humped-shaped relationship between estimated CH_4_ emission rates and inundation time (D). The x-axis in (D) is a proxy to inundation duration of the system as these sites were inundated around the same time. *p*CO_2_(a) is partial pressure of atmospheric CO_2_. Atmospheric δ^13^C_DIC_(eq) represents the δ^13^C of DIC when δ^13^C of 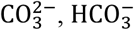, and CO_2(aq)_ is all in equilibrium with δ^13^C of atmospheric CO_2_. Error bar in (C) indicates 95% creditable interval of Bayesian posterior estimates, and error bar in (D) indicates ± standard deviation of estimated CH_4_ emission rates across all sites. We had smaller dataset in January because of no sampling in January, 2022.

### Mineralization and methanogenesis of t-OM as DIC sources

δ^13^C of DIC varied widely over time and space. In the rice field and Wetlands 1 and 2, δ^13^C of DIC enriched from January to early May in both 2021 and 2022 (red, green, and brown lines in Fig. 2B), until it was well above the atmospheric δ^13^C_DIC_(eq) (−0.17 ∼ 0.92 ‰). When δ^13^C of DIC was greater than δ^13^C_DIC_(eq) (e.g., δ^13^C_DIC_ > 2 ‰), it became increasingly depleted in May at Wetland 1, returning to δ^13^C_DIC_(eq), partly driven by air-water CO_2_ exchange. This depletion also occurred at Wetlands 3 and 5 between March and May, 2022 (Fig. 2B). Overall, the seasonal differences in δ^13^C of DIC reached 10.5 ‰ at single site, and 18.24 ‰ cross all sites. Such a high magnitude of variation in δ^13^C of DIC indicates shifts in relative strengths of different DIC-related biogeochemical processes.

DIC and DI^13^C mass balance modeling reveals that both air-water CO_2_ diffusion and methanogenesis likely enriched *δ*^13^C of DIC from January to May (Fig. 2C). When δ^13^C of DIC was well below the atmospheric δ^13^C_DIC_(eq) (e.g., rice field and Wetland 2), air-water diffusion was an important enriching process. The contribution of methanogenesis in the enrichment varied over time. In January, methanogenesis and mineralization had almost equal contributions to DIC, except for Wetland 2 where methanogenesis was lower than mineralization (the site with depleted *δ*^13^C of DIC and slower *δ*^13^C enrichment rate from January to March, Fig. 2B). In contrast, in March, DIC from methanogenesis dominated DIC sources across all sites (Fig. 2C), resulting in enriched *δ*^13^C of DIC and very high CH_4_ emission rates (∼387 mg C m^-2^ d^-1^). However, the emission rates dropped in May, forming a hump-shaped relationship between CH_4_ emission and inundation duration (Fig. 2D). This hump-shaped pattern was not very sensitive to the ratio of CH_4_ oxidized in water to total CH_4_ produced by methanogenesis we assumed in the model (Table S2).

In January-March, estimated aquatic photosynthesis rates were low due to lower phytoplankton concentration (Chla < 20 mg m^-3^; Fig. S5A). In May, aquatic photosynthesis became a large DIC sink at some sites (Wetlands 1 and 3) due to high phytoplankton density occurring with warmer temperature and shallower water (Chla > 120 mg m^-3^; Fig. S5A). Enhanced photosynthesis raised dissolved oxygen (supersaturated), accelerating TOC mineralization in these sites. However, even with large CO_2_ consumption during the high rates of aquatic photosynthesis, these wetlands continued to emit CO_2_. Therefore, most DIC used for aquatic photosynthesis was unlikely from the atmosphere, but from mineralization and methanogenesis of t-OM.

### DIC support of phytoplankton and zooplankton and its spatiotemporal variation

The large variation in *δ*^13^C of DIC allows us to examine how DIC is transferred between primary producers and consumers. We found that *δ*^13^C of phytoplankton varied almost linearly with *δ*^13^C of DIC (slope = 0.85, *R*^2^ = 0.87, *p* < 0.001; Fig. 3A). *δ*^13^C of DOC and POM was not significantly different (*t* = 0.59, *df* = 8, *p* = 0.57) and had relatively small variances (Fig. 3H). DOC and POM, both of which are mixtures from derived phytoplankton and t-OM, represent resources available in water column; however, *δ*^13^C of Daphnia and Copepoda was correlated with *δ*^13^C of DIC with a steeper regression slope than the slope between *δ*^13^C of DOC/POM and *δ*^13^C of DIC, closer to the regression slope between *δ*^13^C of phytoplankton and *δ*^13^C of DIC (Fig. 3A-E). This suggests that zooplankton used a higher proportion of phytoplankton than the proportion of phytoplankton available in water column. Copepoda showed a stronger preference for phytoplankton than did Daphnia, as *δ*^13^C of Copepoda was more correlated to *δ*^13^C of DIC, exhibiting a steeper regression slope and a tighter linear relationship than that of *δ*^13^C of Daphnia (Fig. 3B-C). Daphnia were more flexible in using resources available in the environment, because Daphnia’s *δ*^13^C was more highly correlated with *δ*^13^C of POM than with *δ*^13^C of DIC (Fig. 3C and G).

**Figure 3.**
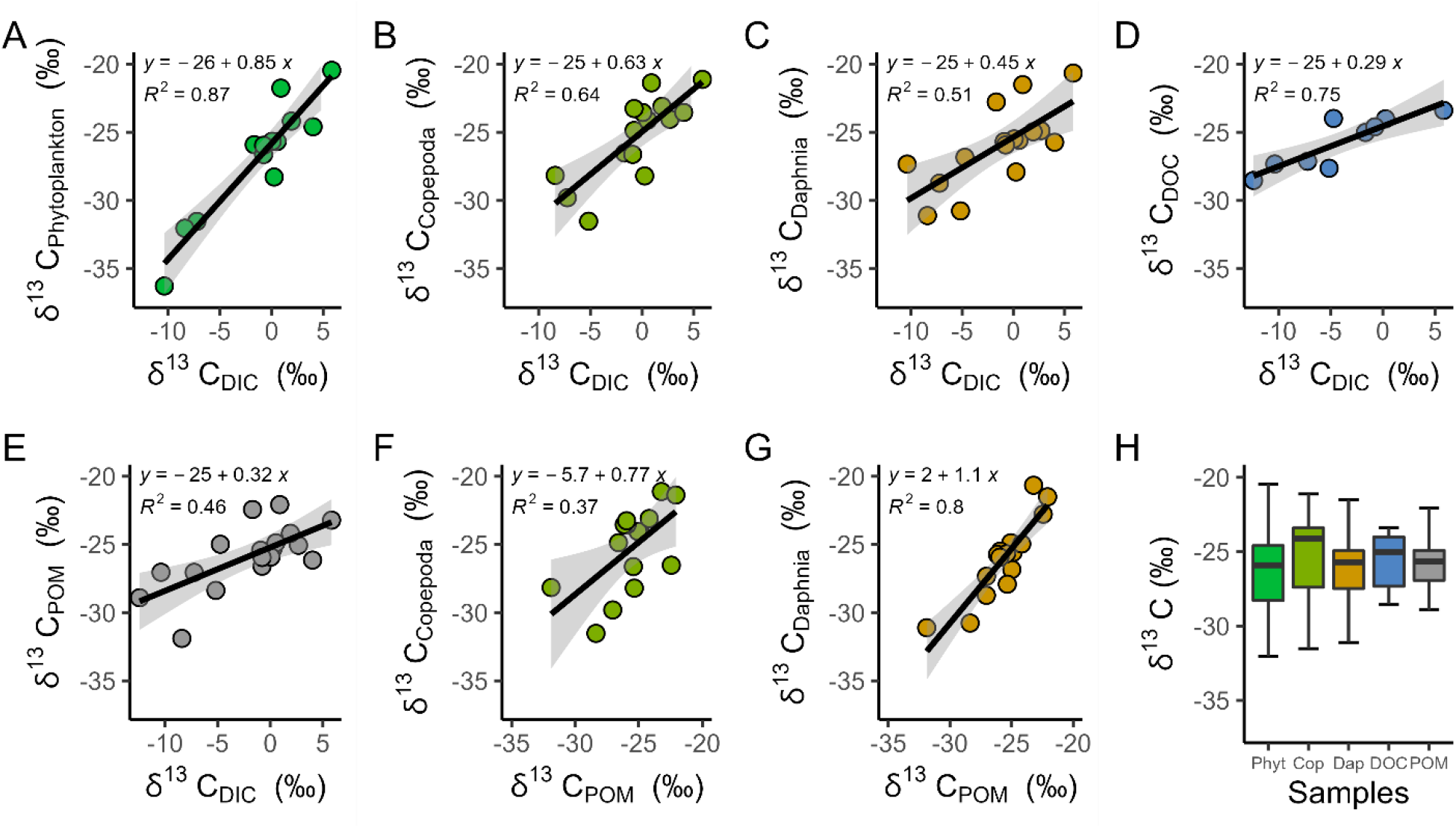
Strong correlations among *δ*^13^*C* of DIC, *δ*^13^*C* of phytoplankton, *δ*^13^*C* of Daphnia, *δ*^13^*C* of Copepoda, *δ*^13^*C* of DOC, and *δ*^13^*C* of POM (A-G), and comparison of *δ*^13^*C* (across sites and all sampling events) in phytoplankton, Copepoda, Daphnia, DOC, and POM (H). *p*-values in (A-G) are all < 0.02.

Our mixing modeling shows that the contribution of phytoplankton to zooplankton biomass shifted over time. In January, when phytoplankton concentrations were low, ∼70 % of Daphnia biomass (Copepoda biomass was low) was supported by t-OM via the OC pathway (Figs. 1B and 4A). In March and May, phytoplankton using DIC from t-OM decomposition constituted ∼68 % of zooplankton biomass (Fig. 4). The contribution of carbon from methanotrophs to zooplankton biomass is minimal (0∼5.8 %), and *δ*^13^C of zooplankton was distinct from the highly depleted *δ*^13^*C* signature of methanotrophs (∼−68 ‰) (Fig. 3H).

**Figure 4.**
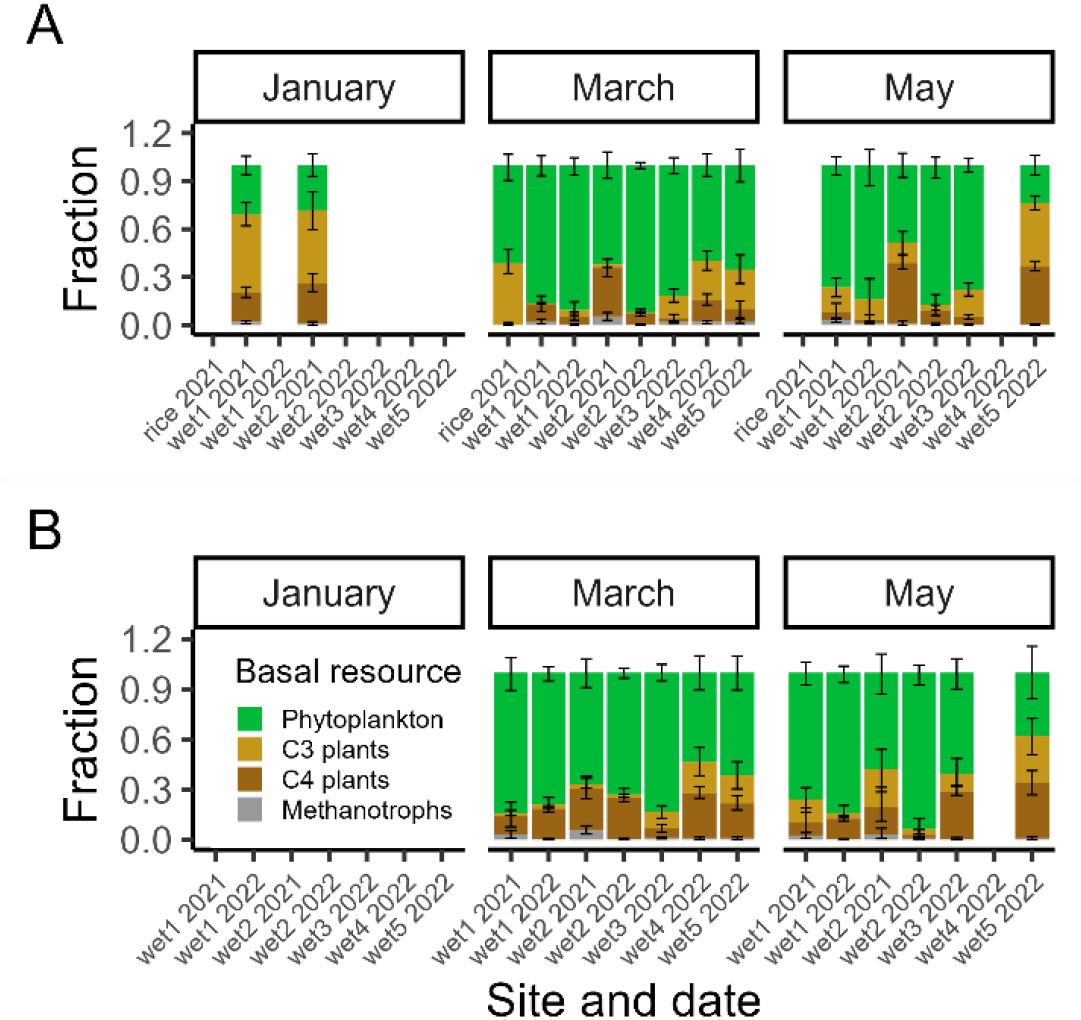
Potential relative contributions of t-OM (C3 and C4 plants), phytoplankton, and methanotrophs to the biomass of Daphnia (A) and Copepoda (B) over time. Note: we did not sample in January, 2022, and Copepoda density in January, 2021 was too low to collect enough samples for stable isotope analysis

## Discussion

Our study provides strong evidence for terrestrial support of aquatic food webs through the DIC pathways. In our system, the varying contribution of methanogenesis to DIC resulted in large variations in *δ*^13^C of DIC (Fig. 2B), comparable to the magnitude of fluctuations of *δ*^13^C of DIC in ^13^C addition experiments (Carpenter et al. 2005; Taipale et al. 2008). Therefore, our system functioned as a natural laboratory to study sources and fates of DIC. We found that in March and May zooplankton primarily fed on phytoplankton (Fig. 4), but these primary producers actually used DIC recycled from t-OM decomposition. This overlooked DIC pathway is likely prevalent in other aquatic ecosystems, particularly these shallow, lentic ecosystems rich in fresh t-OM (Lennon et al. 2006; Demars et al. 2020). The importance of this pathway varied with time. In January when growth of primary producers was limited, probably by low temperature (Fig. S5B), the DIC pathway was weakened, but the OC pathway was strengthened through the flexible diet of Daphnia (Jeffres et al. 2020). The two compensatory pathways originating from t-OM reveal the pivotal role of t-OM in maintaining productivity and stability of aquatic communities (Moore et al. 2004; Nakamoto et al. 2022). The role of CH_4_ pathway was minimal in our system, likely because of low methanotroph density and CH_4_ oxidation rates typical of shallow aquatic ecosystems (Zhang et al. 2020).

Accompanying the DIC pathway, we estimated high CH_4_ emission rates. One of the largest uncertainties in the global CH_4_ budget stems from these under-studied, small, shallow, temporary aquatic ecosystems (Hondula et al. 2021b). Our study partially fills this gap by revealing three features potentially typical for such ecosystems. First, our estimated CH_4_ emission rates were ∼ 1.5-3 times higher than those recorded in nearby wetlands measured by ecosystem-scale eddy covariance flux (38 ∼ 267 mg C m^-2^ d^-1^ vs Fig. 2D) (Anderson et al. 2016; McNicol et al. 2020). This discrepancy is likely because inundated areas have much higher emission rate than non-inundated areas (Hondula et al. 2021b), but measurements by eddy covariance flux capture the mean of inundated and non-inundated areas. Direct field measurements of CH_4_ flux are needed to validate our model-estimated results. Second, we found a hump-shaped relationship between CH_4_ emission rate and inundation duration (Fig. 2D). In addition to the well-recognized effects of high temperature promoting CH_4_ emission (McNicol et al. 2020), hydrologic regime such as duration of inundation may be more critical in regulating CH_4_ emission in seasonal/intermittent aquatic ecosystems (Hondula et al. 2021b). A better estimate of global CH_4_ budget will consider not only varying inundated areas over time (Hondula et al. 2021a), but also varying emission rates with hydrologic regime. Lastly, the wide variation in *δ*^13^C of DIC exhibited in our system might be a feature shared by those small, shallow, temporary aquatic ecosystems with large t-OM inputs. This speculation is partly supported by the increasing variation in *δ*^13^C of DIC with declining lake area (Bade et al. 2004). The high sensitivity of *δ*^13^C of DIC to the shift in biogeochemical processes in shallow aquatic ecosystems makes stable isotope analyses, combined with gas flux measurements, powerful tools to quantify CO_2_ and CH_4_ emission, to identify various sources and sinks of DIC, and to track energy transfer in food webs.

Although wetlands have long been considered to be global carbon sinks (Nahlik and Fennessy 2016), we show that inundated components can be large carbon sources due to abundant t-OM inputs, offsetting carbon sequestration capacity of wetland plants. Likewise, many freshwater ecosystems receive substantial t-OM inputs from their terrestrial surroundings (Cole et al. 2007). In such systems, a large proportion of DIC consumed by aquatic primary producers might be from DIC recycled from t-OM, during which a great amount of CO_2_ and CH_4_ emission occurs, significantly reducing terrestrial carbon sinks (Webb et al. 2019). Focusing solely on the OC pathways between terrestrial and aquatic ecosystems might miss the big picture of multiple pathways by which t-OM can support aquatic ecosystems (Fig. 1B), and underestimate the role of freshwater ecosystems in the global carbon cycle by storing, respiring, transferring, and transporting t-OM (Liu et al. 2022). With widespread warming effects of global climate change, such small, shallow, temporary aquatic ecosystems might become more prevalent as perennial streams turn into intermittent ones, permanent wetlands become seasonal, and intense floods intermittently submerge additional lands (Hammond et al. 2021). As ecosystems alternate between aquatic and terrestrial states, they may exhibit new CO_2_ and CH_4_ emission patterns, thereby creating feedbacks that potentially exacerbate (or mitigate) effects of global climate change.

## Acknowledgements

We thank Carson Jeffres for his inspiring discussion of this work. Thanks to Jocelyn Rodriguez for her assistance in collecting field samples. We appreciate the help of Steven Sadro for generously allowing us to use his laboratory and equipment. This research was partially funded by a Yolo Basin Foundation Graduate Student Fellowship to JW. We acknowledge the support from ASLO and Wiley in the form of an APC waiver offered to JW via the L&O Letters Early Career Publication Honor.

## Supplementary information

### Text S1 Details about sample collection, processing, and measurement

We conducted six field sampling events in total. In each sampling event, we collected water samples, surface sediments, phytoplankton (i.e., algae floating on water surface, in water column, or associated with plants), zooplankton, and terrestrial detritus at 2∼3 replicated locations with water depth <0.3 m. After collection, these samples were immediately placed in ice water. We measured water depth and in-situ water quality including temperature, pH, Chlorophyll-a (Chla), and dissolved oxygen using a handheld YSI EXO2 multiparameter Sonde. The Sonde measured Chla was calibrated with lab measured Chla in 2021. Each water sample was used for four purposes. We sent 200 ml water samples to the analytical lab at the University of California, Davis (https://anlab.ucdavis.edu/) to measure concentrations of DIC, DOC, and TOC. We filtered remaining water samples through three glass fiber filters with 0.7 *μm* pore size (Millipore, AP4002500) until each filter reached impermeability. We sent 30 ml filtered water to measure *δ*^13^*C* of DOC. Another fraction of filtered water was used to fill a 10 ml glass vial completely (i.e., have a positive meniscus) to measure *δ*^13^*C* of DIC. Filters with remaining seston were used to measure *δ*^13^*C* of POM. In the lab, we washed all the solid samples except sediments with DI water, dried them for 48 hours under oven temperature of ∼ 60 °C, and ground them. Dried sediment samples were further acid fumigated to remove carbonate using concentrated HCl (12 M) and encapsulated in silver capsules. Lastly, all dried solid samples were encapsulated in tin capsules to measure *δ*^13^*C* and *δ*^15^*N* in solids. All the stable isotope analyses were conducted at the University of California, Davis Stable Isotope Facility using isotope ratio mass spectrometer (https://stableisotopefacility.ucdavis.edu/).

It is worth noting that our study sites dried at different times, which caused no data at some sites in May. The rice field was drained by farmers in early March to grow rice, and the Wetland 4 dried naturally in April, earlier than other wetlands. These resulted in no data in May for the rice field or Wetland 4 (Fig. 2). Most wetlands (Wetlands 1, 3, and 5) dried naturally in May, except the relatively larger and deeper Wetland 2, which dried in June. The maximum water surface area of these wetlands varies between 0.23∼0.62 km^2^ across the six sites.

### Text S2 Method to determine variables and parameters in the DIC and DI^13^C mass balance model

The DIC and DI^13^C mass balance models (Eqs. 1-2) include various variables and parameters that can be classified into four categories. The first category is the variables that are directly measured, including variables describing concentrations of water chemistry (i.e., [DIC], [TOC], and [Chla]), water depth (*z*), and partial pressure of CO_2_ in the air (*pCO*_2_(*a*)), which is ∼419 ppm during the sampling period (https://scrippsco2.ucsd.edu/).

The second category is derived variables, including aqueous CO_2_ concentration ([CO_2_(aq)]), [*DI* ^13^*C*], and ^13^*C*/^12^*C* ratios (i.e., 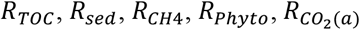, and 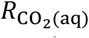). [CO_2_(aq)] was calculated based on measured DIC concentration ([DIC]), pH, and water temperature by assuming that the three DIC species, i.e., CO_2_(aq), 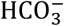, and 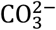, are at equilibrium. Thus, [CO_2_(aq)] was derived by solving the following mass balance and chemical equilibrium equations (Chapra et al. 2008).

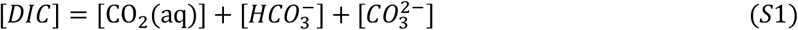

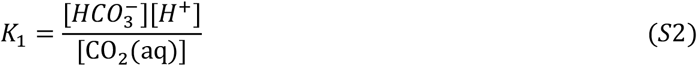

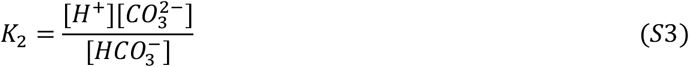

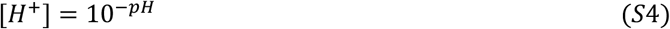

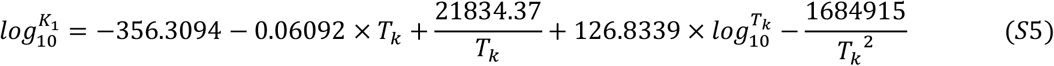

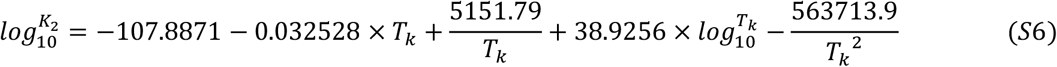

where 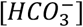 and 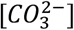 are concentrations of bicarbonate and carbonate (mol m^-3^); [*H*^+^] is hydrogen-ion concentration (mol L^-1^); *K*_1_ and *K*_2_ are the first and second acidity constants (mol L^-1^); and *T*_*k*_ is Kelvin temperature (K).

We calculated ^13^*C*/^12^*C* ratios of samples containing carbon (i.e., 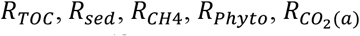, and 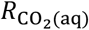) based on their measured or derived *δ*^13^*C* using the definition of *δ*^13^*C* as follows:

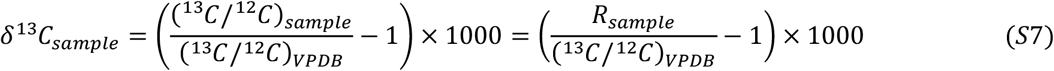

where (^13^*C*/^12^*C*)_*VPDB*_ is the ^13^*C*/^12^*C* ratio of Vienna Pee Dee Belemnite, which is used to standardize ^13^*C*/^12^*C* ratio of a sample. We measured *δ*^13^*C* in sediment organic carbon (*δ*^13^*C*_*sed*_) and in phytoplankton (*δ*^13^*C*_*phyto*_). We did not measure *δ* ^13^*C* of methane (*δ*^13^*C*_*CH*4_) directly; instead, we used existing data of *δ*^13^*C*_*CH*4_ (−68.87 ± 1.76 ‰) reported for wetlands in California, measured close to (∼ 55 km) our study sites (McNicol et al. 2020). We obtained *δ*^13^*C* of CO_2_ in the air 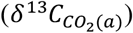 from the Scripps CO_2_ program (https://scrippsco2.ucsd.edu/). We derived *δ*^13^*C* of TOC (*δ* ^13^*C*_*TOC*_) based on measured *δ*^13^*C* of DOC (*δ*^13^*C*_*DOC*_), *δ*^13^*C* of POC (*δ*^13^*C*_*POC*_), and measured concentrations of DOC and POC using weighted average of *δ* ^13^*C*_*DOC*_ and *δ* ^13^*C*_*POC*_, described as

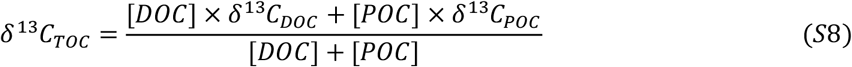

We derived *δ*^13^*C* of aqueous 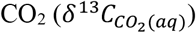 based on measured *δ*^13^*C* of DIC (*δ* ^13^*C*_*DIC*_), derived concentrations of the three DIC species (see Eqs. S1-S6), and the equilibrium fractionation effects between aqueous CO_2_, gaseous CO_2_, bicarbonate, and carbonate following the approach proposed by Zhang et al. (1995), Venkiteswaran et al. (2014), and Campeau et al. (2017). This approach is described as follows:

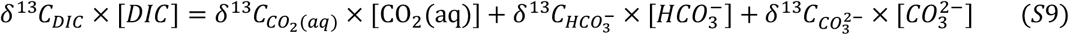

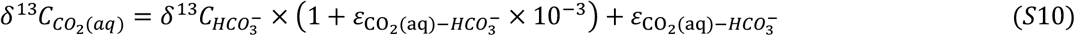

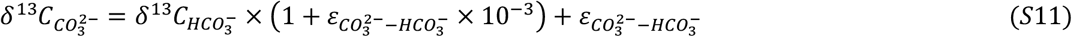

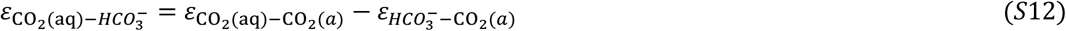

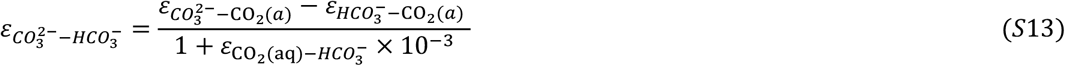

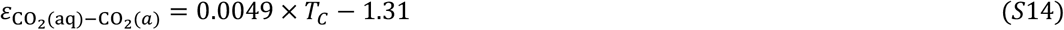

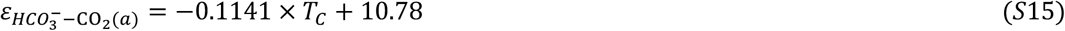

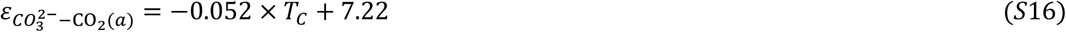

where 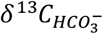 and 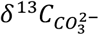 are *δ*^13^*C* signatures of bicarbonate and carbonate (‰); 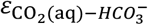 is the permil equilibrium fractionation between aqueous CO_2_ and bicarbonate (‰); 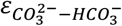 is the permil equilibrium fractionation between carbonate and bicarbonate (‰); 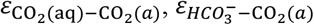, and 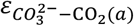 are the permil equilibrium fractionations between aqueous CO_2_ and gaseous CO_2_, between bicarbonate and gaseous CO_2_, and between carbonate and gaseous CO_2_ (‰); and *T*_*C*_ is water temperature (°C). By solving the system of equations (Eqs. S9-S16), we obtained 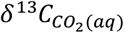 corresponding to each measured *δ*^13^*C*_*DIC*_. Lastly, [DI^13^C] was calculated based on [DIC] and *δ*^13^*C*_*DIC*_ using the following equation:

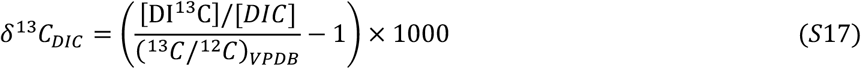

We calculated the time derivative of [DIC] and [DI^13^C] (i.e., 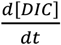 and 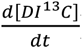) based on measured [DIC] and derived [DI^13^C]. For each site, we used forward difference to calculate the time derivative of one year’s first sampling event, backward difference to calculate the time derivative of one year’s last sampling event, and central difference to calculate the time derivative of one year’s the other sampling events. If one site only has one sampling event for a year (e.g., Wetland 4 in 2022), we could not directly calculate the time derivative and assumed the time derivative equal to 0.

The third category is known parameters, including gas exchange coefficient of CO_2_ (*ω*) and that of ^13^CO_2_ (^13^*ω*), solubility (i.e., Henry constant) of CO_2_ (*β*) and solubility of ^13^CO_2_ (^13^*β*). According to previous studies (Himmelblau 1964; Edmond and Gieskes 1970), the four parameters are all a function of water temperature, given by

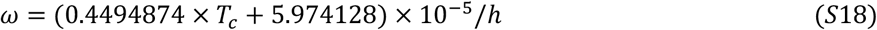

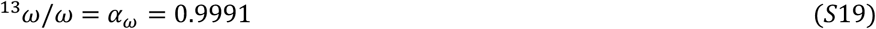

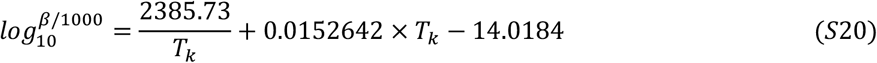

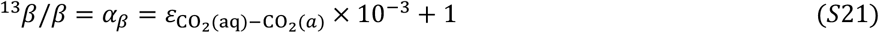

where *h* is stagnant film thickness (m), varying between 0.00027 and 0.0008 m in ocean water; we set *h* = 0.00035 m according to Wanninkhof (1985) and Bade et al. (2004); *α*_*ω*_ is the kinetic fractionation factor during CO_2_ transfer at the air-water interface (Wanninkhof 1985) (unitless); and *α*_*β*_ is the ratio of solubilities of ^13^CO_2_ and CO_2_, which is equal to the equilibrium fractionation between aqueous and gaseous CO_2_ (Zhang et al. 1995).

The last category is unknown parameters in the mass balance equations of DIC and DI^13^C, including *k*_*TOC*_, *MT*_*sed*_, and *k*_*photosyn*_, under the assumption that the oxidized fraction of total CH_4_ produced by sediment methanogenesis is 30% (i.e., *OX*_*CH*4_ = 0.3 × *MT*_*sed*_ /*z*) (McNicol et al. 2020; Zhang et al. 2020). Here, we have three unknowns with two equations for each sampling event at each site. To reduce the number of parameters, we assumed that *k*_*photosyn*_ is constant over all the study sites and sampling events. The implication of this assumption is that we assume aquatic photosynthesis rate is proportional to Chla concentration. With this assumption, we in total has 2×*n* equations and 2×*n*+1 parameters, where *n* is the total number of samplings we conducted for all the sites (*n* = 18, Table S1). We inferred the mean value of each unknown parameter with 95% credible interval using Bayesian Markov chain Monte Carlo method, implemented by the R package *Rstan* (Carpenter et al. 2017).

After estimating the unknown parameters, we calculated each term in Eq. (1) for every sampling event at each site. The CO_2_ emission rate is represented by the last term (*ω* × (β × *pCO*_2_(*a*) − [CO_2_(aq)]). Methane emission rate is represented by 0.7 × *MT*_*sed*_, that is, 30% of produced methane was oxidized and the remaining 70% was emitted.

**Table S1.** Partial pressures of CO_2_ in water column (*p*CO_2(aq)_), their corresponding super-saturation levels, and CO_2_ emission rates in aquatic ecosystems of the rice field and seasonal wetlands.

| Site | Sampling date | $p\text{CO}_{2(\text{aq})}$<br>($\mu\text{atm}$) | Super-saturation level<br>(%) | CO<sub>2</sub> emission<br>($\text{gC m}^{-2} \text{d}^{-1}$) |
| --- | --- | --- | --- | --- |
| Rice field | 2021/01/23 | $4753 \pm 70$ | $1135 \pm 17$ | 0.824 |
| | 2021/03/8 | $9023 \pm 552$ | $2153 \pm 132$ | 1.678 |
| Wetland 1 | 2021/01/23 | $3002 \pm 6$ | $717 \pm 1$ | 0.491 |
| | 2021/03/21 | $3261 \pm 171$ | $778 \pm 41$ | 0.554 |
| | 2021/05/02 | $1039 \pm 121$ | $248 \pm 29$ | 0.125 |
| | 2021/05/22 | $1366 \pm 50$ | $326 \pm 12$ | 0.189 |
| | 2022/03/21 | $3157 \pm 150$ | $753 \pm 36$ | 0.544 |
| | 2022/05/01 | $969 \pm 18$ | $231 \pm 4$ | 0.110 |
| Wetland 2 | 2021/01/23 | $2787 \pm 18$ | $665 \pm 4$ | 0.450 |
| | 2021/03/21 | $3198 \pm 47$ | $763 \pm 11$ | 0.542 |
| | 2021/05/02 | $2651 \pm 5$ | $633 \pm 1$ | 0.450 |
| | 2022/03/21 | $3559 \pm 341$ | $849 \pm 81$ | 0.624 |
| | 2022/05/01 | $2193 \pm 5$ | $523 \pm 1$ | 0.356 |
| Wetland 3 | 2022/03/21 | $2496 \pm 496$ | $596 \pm 18$ | 0.413 |
| | 2022/05/01 | $1463 \pm 6$ | $349 \pm 1$ | 0.210 |
| Wetland 4 | 2022/03/21 | $715 \pm 76$ | $171 \pm 18$ | 0.059 |
| Wetland 5 | 2022/03/21 | $1040 \pm 144$ | $248 \pm 34$ | 0.123 |
| | 2022/05/01 | $1000 \pm 16$ | $239 \pm 4$ | 0.117 |

Note: the number before ' ± ' represents mean value and that after ' ± ' is standard deviation. Super-saturation level is calculated using $p\text{CO}_2$ in water column divided by $p\text{CO}_2$ in the atmosphere ( $\sim 419 \mu\text{atm}$ during the sampling period).

**Table S2.** The sensitivity of estimated CH_4_ emission rates to the oxidized fraction of total CH_4_ produced by sediment methanogenesis (F_CH4_ox_).

| Site | Sampling date | CH<sub>4</sub> emission (mg C m<sup>-2</sup> d<sup>-1</sup>) if F<sub>CH<sub>4</sub>_ox</sub> = 0.1 | CH<sub>4</sub> emission (mg C m<sup>-2</sup> d<sup>-1</sup>) if F<sub>CH<sub>4</sub>_ox</sub> = 0.2 | CH<sub>4</sub> emission (mg C m<sup>-2</sup> d<sup>-1</sup>) if F<sub>CH<sub>4</sub>_ox</sub> = 0.3 | CH<sub>4</sub> emission (mg C m<sup>-2</sup> d<sup>-1</sup>) if F<sub>CH<sub>4</sub>_ox</sub> = 0.4 | Sensitivity ratio |
| --- | --- | --- | --- | --- | --- | --- |
| Rice field | 2021/01/23 | 335.9236 | 338.7419 | 345.8639 | 348.3851 | 0.0364 |
|  | 2021/03/8 | 629.2486 | 646.0238 | 669.1598 | 698.9391 | 0.1055 |
| Wetland 1 | 2021/01/23 | 126.5706 | 131.0826 | 139.4961 | 144.5183 | 0.1325 |
|  | 2021/03/21 | 410.0919 | 430.0784 | 448.3508 | 436.3588 | 0.0887 |
|  | 2021/05/02 | 129.2238 | 142.885 | 169.2222 | 223.4181 | 0.5668 |
|  | 2021/05/22 | 108.7804 | 110.311 | 91.51822 | 33.06876 | 0.8990 |
|  | 2022/03/21 | 424.3042 | 443.0634 | 457.8675 | 392.3363 | 0.1526 |
|  | 2022/05/01 | 213.2612 | 224.296 | 226.0532 | 205.2692 | 0.0957 |
| Wetland 2 | 2021/01/23 | 74.10234 | 74.89686 | 75.19967 | 78.49859 | 0.0581 |
|  | 2021/03/21 | 439.7843 | 454.9942 | 473.3672 | 465.6488 | 0.0733 |
|  | 2021/05/02 | 308.746 | 321.5734 | 332.3835 | 326.9012 | 0.0733 |
|  | 2022/03/21 | 505.9064 | 502.7101 | 498.2906 | 467.5344 | 0.0777 |
|  | 2022/05/01 | 407.5396 | 424.775 | 437.7426 | 476.7211 | 0.1584 |
| Wetland 3 | 2022/03/21 | 261.7926 | 272.4568 | 286.171 | 266.1687 | 0.0897 |
|  | 2022/05/01 | 127.8718 | 132.8766 | 138.7931 | 149.6488 | 0.1586 |
| Wetland 4 | 2022/03/21 | 46.52734 | 42.44963 | 40.56601 | 50.66953 | 0.2243 |
| Wetland 5 | 2022/03/21 | 235.5038 | 241.2797 | 223.8354 | 220.76 | 0.0891 |
|  | 2022/05/01 | 161.5306 | 172.8547 | 185.9613 | 205.6026 | 0.2428 |

Note: we defined sensitivity ratio as the range of estimated CH<sub>4</sub> emission rates / the average of estimated CH<sub>4</sub> emission rates, when F<sub>CH<sub>4</sub>\_ox</sub> was varied between 0.1 and 0.4 (i.e., the oxidized fraction range observed in shallow aquatic ecosystems) (Martinez-Cruz et al. 2015; McNicol et al. 2020; Zhang et al. 2020). This sensitivity analysis shows that estimated methane emission rates are much sensitive to the oxidized fraction only at Wetland 1 during May, 2021 when CH<sub>4</sub> emission rates were relatively low (33-224 mg C m<sup>-2</sup> d<sup>-1</sup>).

**Figure S1.**
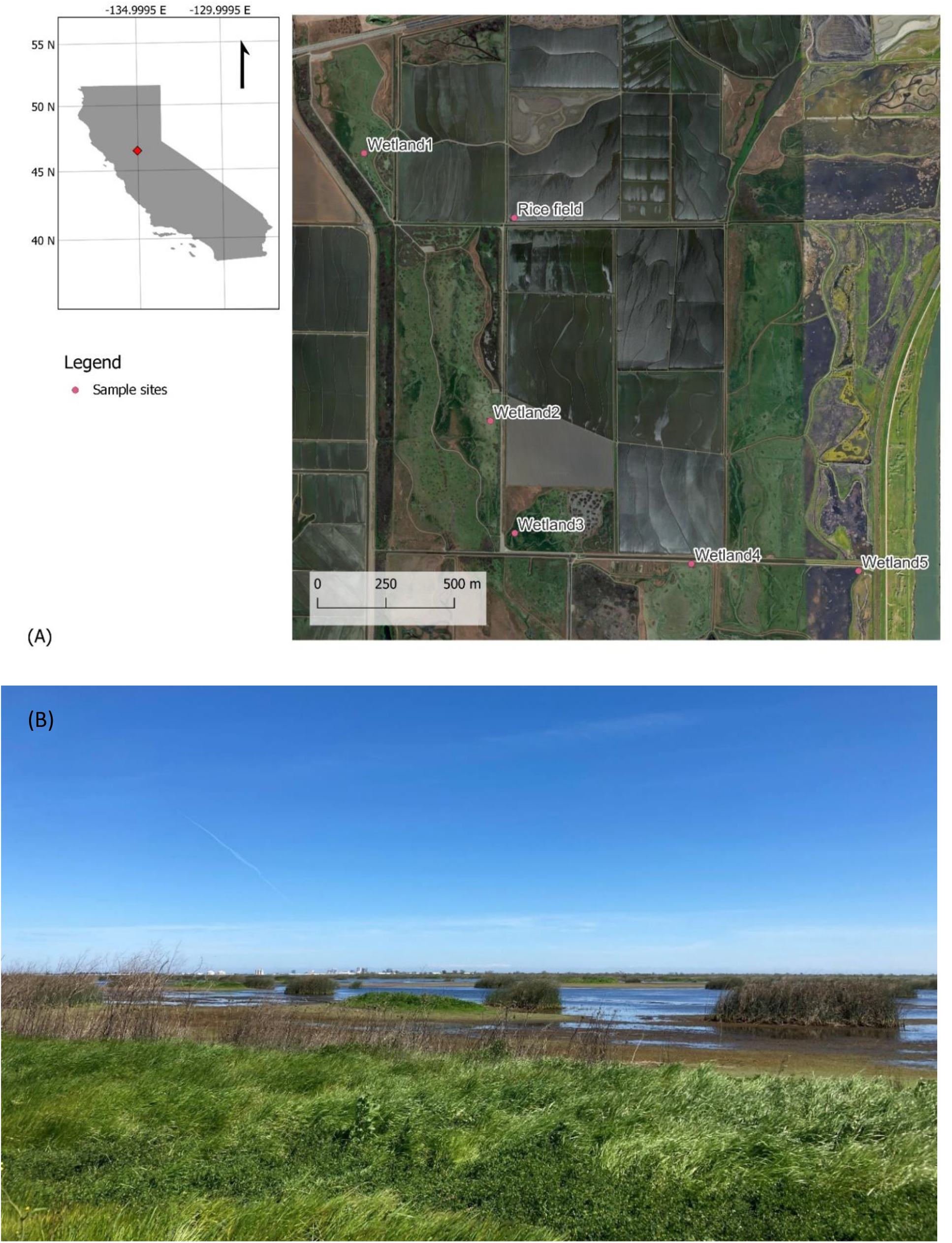
Geographic locations of our study sites (121.62W, 38.54N) in the Yolo Bypass Wildlife area, California (A) and a photo of one study site Wetland 1 (B). As shown in this photo, during the inundated season, emergent plants mostly grow on islands and edges of these wetlands.

**Figure S2.**
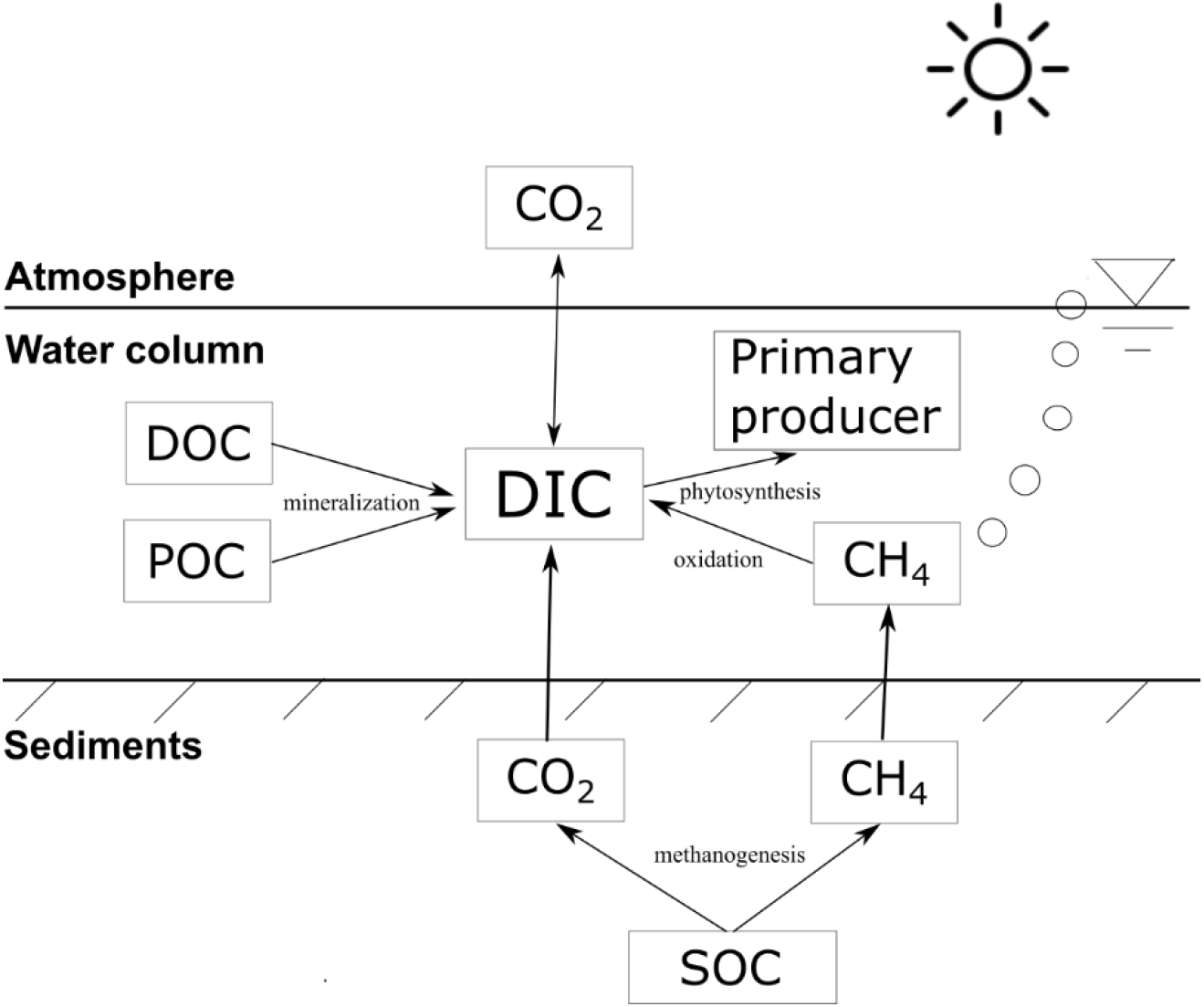
Conceptual diagram of dissolved inorganic carbon (DIC) sources and sinks in aquatic ecosystems of seasonal wetlands. DOC is dissolved organic carbon, POC is suspended particulate organic carbon, and SOC is sediment organic carbon.

**Figure S3.**
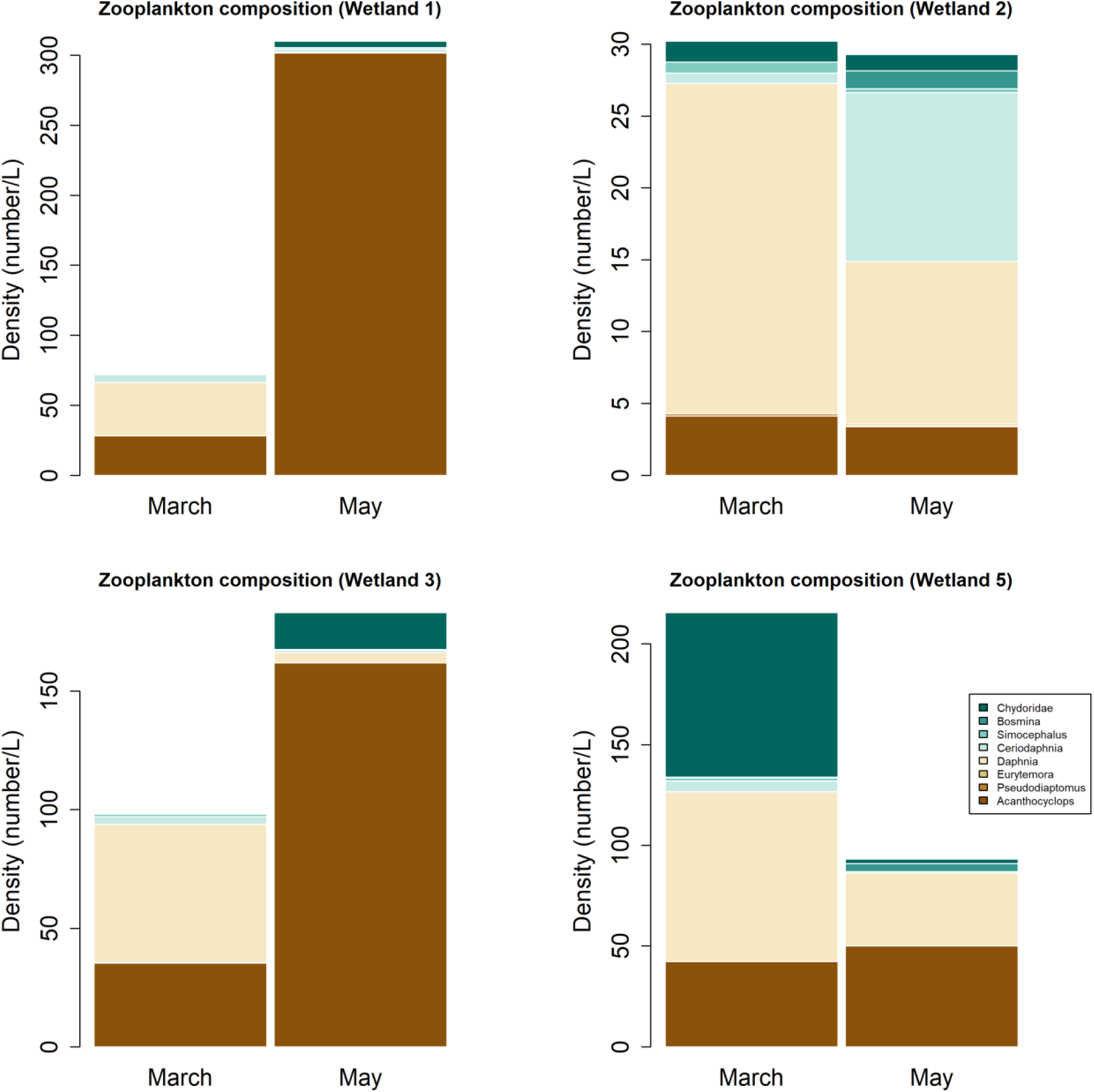
Seasonal variation in density and composition of zooplankton across four wetland sites (Wetlands 1-3 and 5) in 2022. We did not show the data in Wetland 4 because it was dried in April, and the data in March are quite similar to Wetland 5. Here, Copepoda include *Acanthocyclops, Pseudodiaptomus*, and *Eurytemora*. The other five zooplankton taxa belong to the order Cladocera. Although Chydoridae is more abundant than Daphnia in some cases (May at wetland 3), body size of Daphnia is ∼ 7 times higher than that of Chydoridae by surface area.

**Figure S4.**
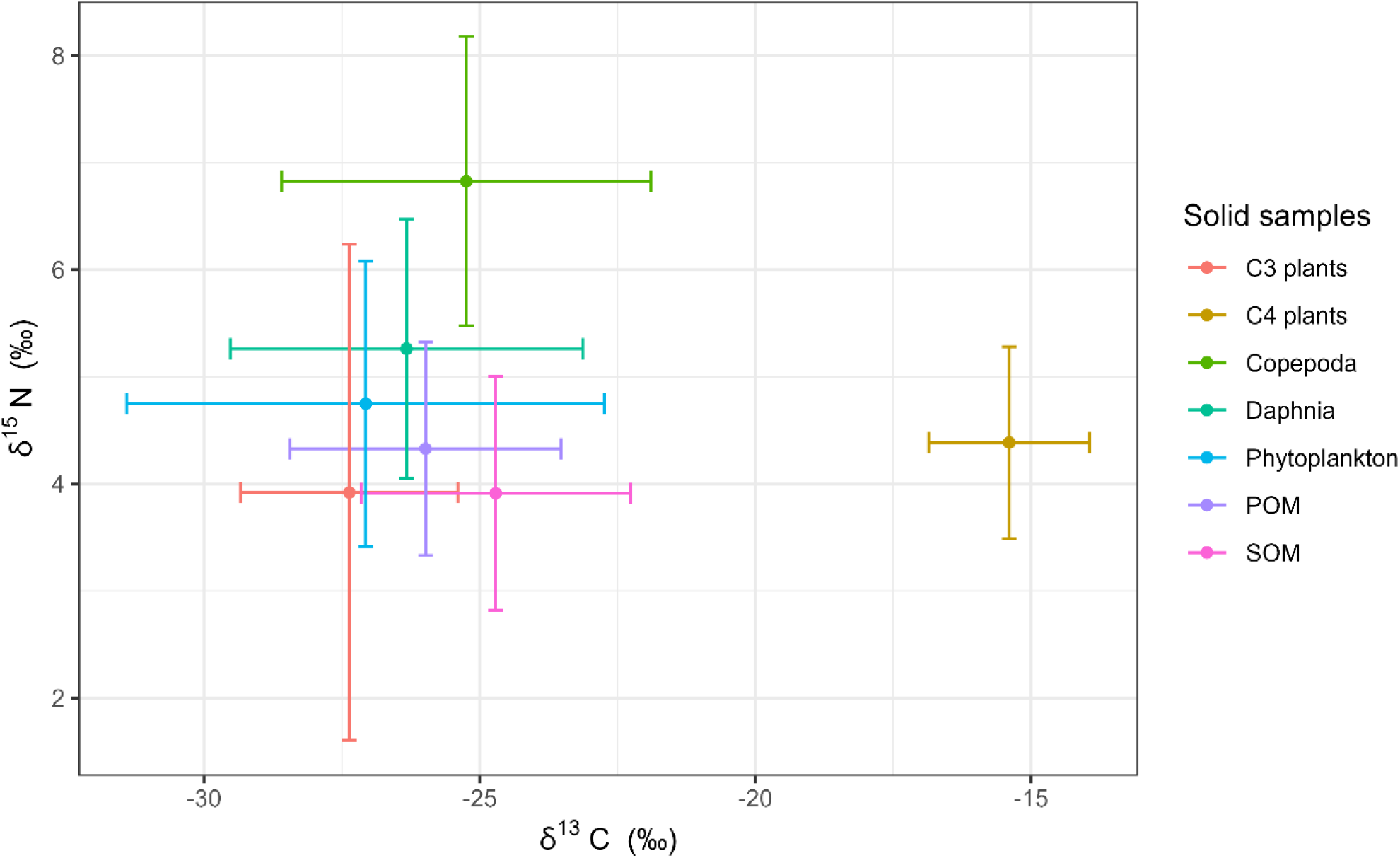
*δ*^13^*C* and *δ*^15^*N* signatures (means and standard deviations) of various solid samples collected from all the sites through all sampling events.

**Figure S5.**
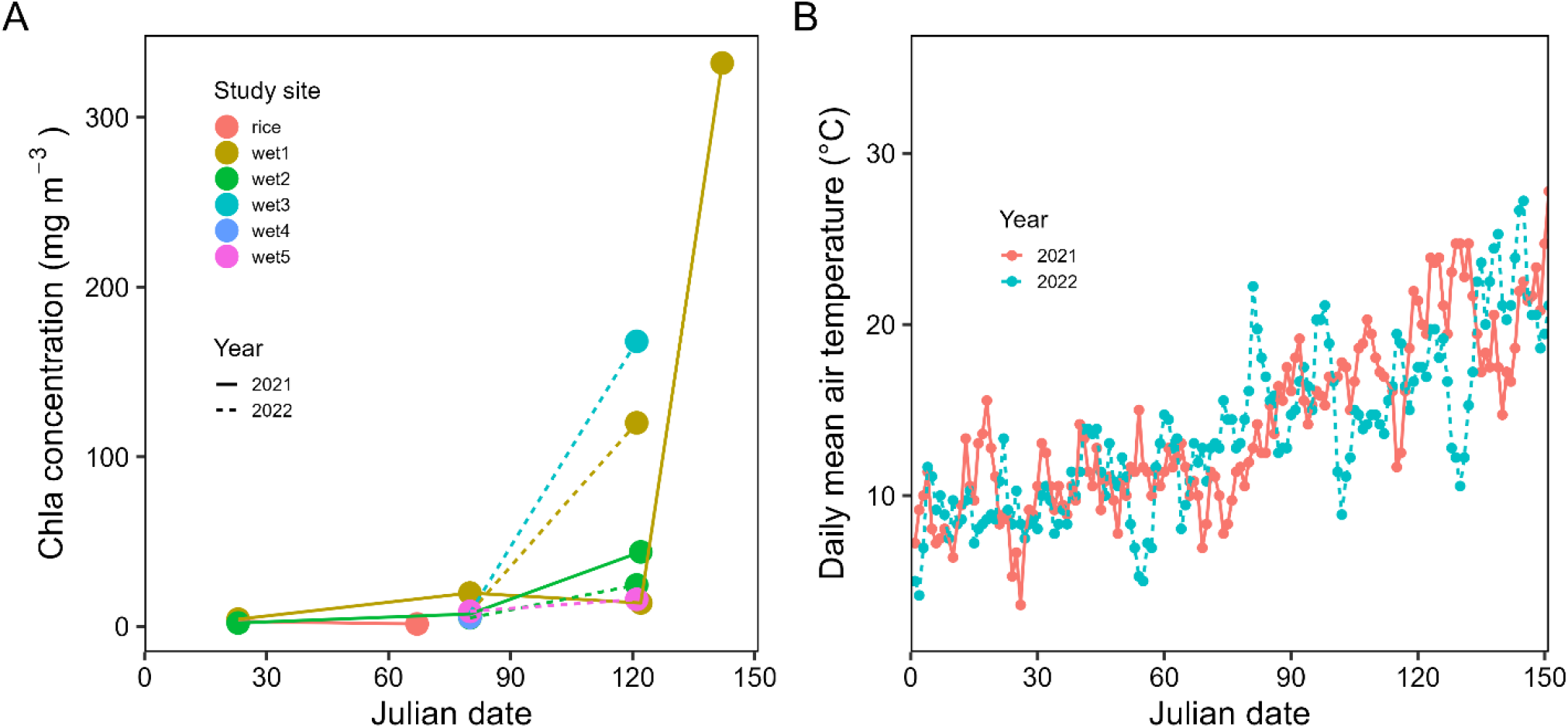
Temporal variations in chlorophyll-a (Chla) across the six study sites (A) and in daily mean air temperature (B) at west Sacramento (∼10 km from our sites).

